# Vaginal Isolates of *Candida glabrata* are Uniquely Susceptible to Ionophoric Killer Toxins Produced by *Saccharomyces cerevisiae*

**DOI:** 10.1101/2020.10.14.339788

**Authors:** Lance R. Fredericks, Mark D. Lee, Hannah R. Eckert, Shunji Li, Mason A. Shipley, Cooper R. Roslund, Dina A. Boikov, Emily A. Kizer, Jack D. Sobel, Paul A. Rowley

**Author notes:** Authors contributed equally (Hannah R. Eckert, Shunji Li, Mason A. Shipley, Cooper R. Roslund).

## Abstract

Compared to other species of *Candida* yeasts, the growth of *Candida glabrata* was inhibited by many different strains of *Saccharomyces* killer yeasts. The ionophoric K1 and K2 killer toxins were broadly inhibitory to all clinical isolates of *C. glabrata* from patients with recurrent vulvovaginal candidiasis, despite high levels of resistance to clinically relevant antifungal therapeutics.

## RESEARCH FINDINGS

Vulvovaginal candidiasis (VVC) is estimated to afflict two in every three women worldwide at some point in their lives, causing significant suffering and associated economic losses (1–3). *Candida albicans* is most often isolated as the dominant species present in cases of VVC, followed by *Candida glabrata* (4). In certain diabetic patient populations, *C. glabrata* can be the dominant yeast species associated with VVC (5, 6). The main treatment for VVC is the orally administered fungistatic azole, fluconazole. During pregnancy, to relieve symptoms of VVC and to prevent *Candida-*associated complications, topical application of azoles is preferred over oral administration due to the potential for fetal toxicity in the first trimester (7). However, drug-resistance in isolates of *C. glabrata* and other species of *Candida* yeasts is increasing and can result in long courses of suppressive treatment and treatment failure (8–10). The limited availability of effective non-toxic therapies to treat VVC warrants the exploration of novel therapeutics that are active at the normal low pH of the vagina.

Killer toxins produced by *Saccharomyces* “killer” yeasts are optimally active in acidic conditions (pH ≤4.6) that overlap the pH of the vaginal mucosa (~pH 4.2) (11). In addition, there have been many studies describing killer yeasts that can inhibit the growth of pathogenic fungi (12–17). Given the discovery of novel killer toxins produced by *Saccharomyces* killer yeasts, 16 species of the *Candida* genus were screened for their susceptibility to nine killer yeast strains known to express K1, K1L, K2, K21/K66, K28, K45, K62, K74, or Klus killer toxins encoded by double-stranded satellite RNAs (18–24). To test the ability of different killer yeasts to inhibit the growth of *Candida* yeasts, a well assay was used to inoculate killer yeasts into killer assay agar plates (YPD agar plates with 0.003% w/v methylene blue buffered to pH 4.6 with sodium citrate (25)) seeded with ~1 × 10^5^ *Candida* yeast cells (Fig. S1). After three days of incubation at room temperature, growth inhibition was identified by the appearance of yeast-free zones and/or halos of oxidized methylene blue around killer yeasts (Fig. 1 and S1). Of the 16 species of *Candida* yeasts challenged, *C. glabrata* appeared to be the most susceptible to killer yeasts and was only resistant to the killer yeast that expressed K62 (Fig. S1 and Table S1). One additional species of the *Nakaseomyces* genus (*C. nivariensis*) was also more susceptible to *Saccharomyces* killer yeasts than other *Candida* yeasts (Fig. 1A and B). The K21 killer yeast appeared to have the broadest spectrum of antifungal activity (Fig. 1C).

**Fig. 1.**
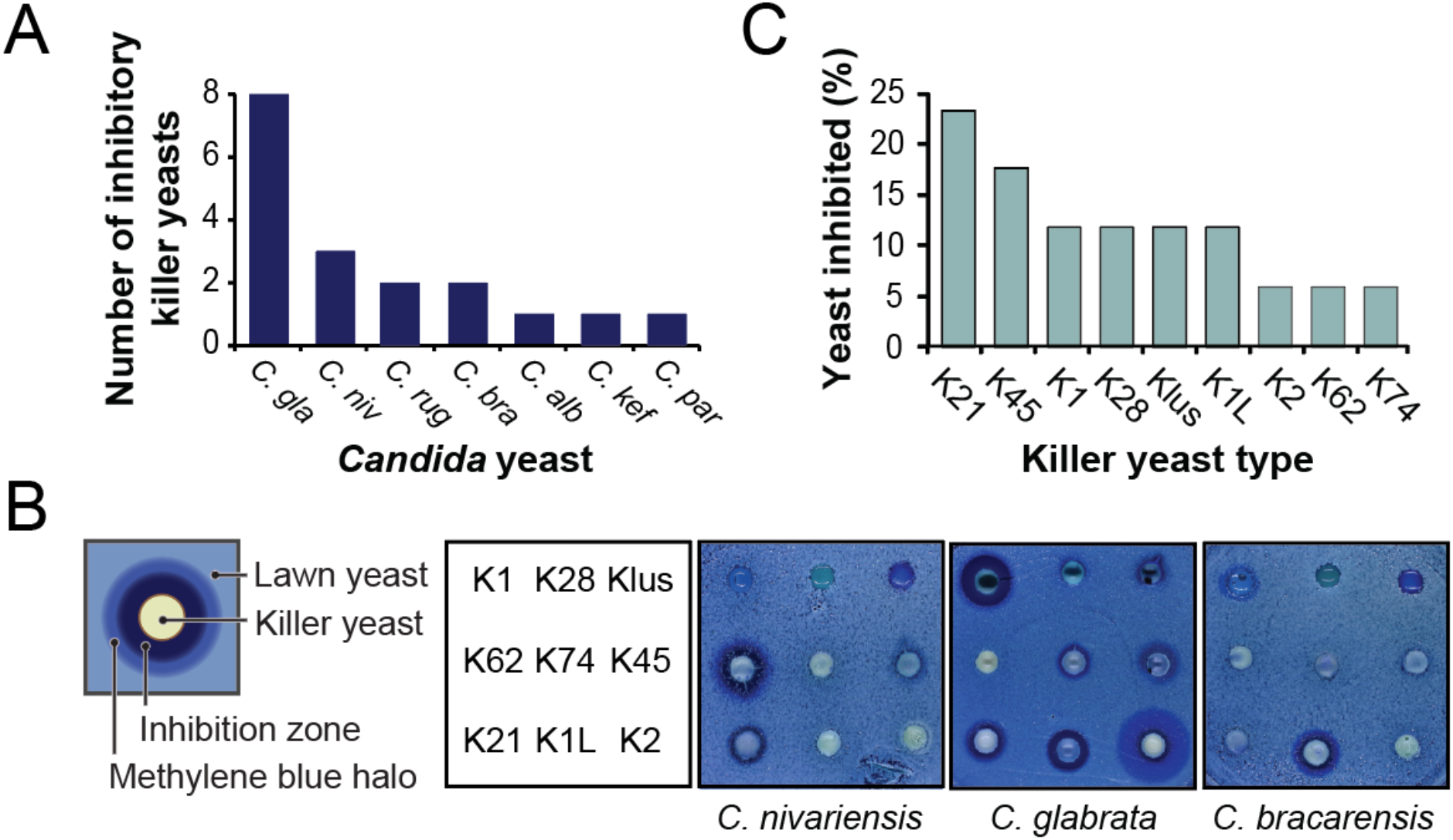
*Candida glabrata* is more susceptible to inhibition by killer yeasts than other species of *Candida* yeasts. (A) The number of killer yeasts that inhibit the growth of different species of *Candida* yeasts; *C. glabrata* (*C. gla*)*, C. nivariensis* (*C. niv*), *C. kefyr* (*C. kef*), *C. bracarensis* (*C. bra*), *C. rugosa* (*C. rug*), *C. pararugosa* (*C. par*), and *C. albicans* (*C. alb*) (n = 2). (B) A schematic illustration of the effect of a killer yeast on the growth of a competing lawn of yeast. Representative well assay plates with nine different killer yeasts on agar seeded with representative species of *Candida* yeasts. (C) The percentage of *Candida* yeast species found to be inhibited by each type of killer toxin (n = 2).

As killer toxin susceptibility can vary widely within a species, 53 unselected clinical isolates of *C. glabrata* from the human vagina were challenged by killer yeasts. These clinical isolates were collected at the Wayne State University Vulvovaginitis clinic in Detroit, MI, between 2015 and 2019. Killer yeasts were inoculated onto the surface of killer assay agar plates seeded with lawns of *C. glabrata* and qualitatively assayed for evidence of growth inhibition. Of the 477 interactions measured between killer yeasts and *C. glabrata*, K1, K2, and K45 killer yeasts inhibited the growth of 100%, 96%, and 75% of *C. glabrata* isolates, respectively (Fig. 2A, top). The remaining killer yeasts each inhibited less than 33% of *C. glabrata* isolates. The susceptibility of *C. glabrata* to K1 and K2 killer yeasts greatly contrasts the widespread resistance of *Saccharomyces* yeasts (Fig. 2A, bottom). To test the susceptibility of the clinical isolates of *C. glabrata* with acute killer toxin exposure, K1 and K2 toxins were partially purified from 1 mL of spent culture medium, as described previously (23). Exposure of lawns seeded with ~1 × 10^5^ of *C. glabrata* cells to K1 or K2 demonstrated a concentration-dependent growth inhibition of *C. glabrata* (Fig. 2B). All isolates of *C. glabrata* were inhibited by K1 and K2 at the highest concentration tested, with K2 exposure resulting in large halos of methylene blue (185.68 mm^2^ ± 28.66; 95% CI) whereas K1 produced larger zones of growth inhibition (92.34 mm^2^ ±11.24; 95% CI) (Fig. 2B). The inhibition of *C. glabrata* is similar to the fungicidal effects of K1 and K2 toxins against susceptible strains of *S. cerevisiae* (Fig. S2). Neither K1 nor K2 were acutely cytotoxic to cultured human cells at physiological pH as measured by an alamar blue viability assay (Fig. S3). Both K1 and K2 retained measurable activity against yeast cells after a 1 hour incubation with human epithelial cells (HeLa cell line) in Dulbecco’s modified eagle medium with 10% serum at pH 7 (Fig. S3).

**Fig. 2.**
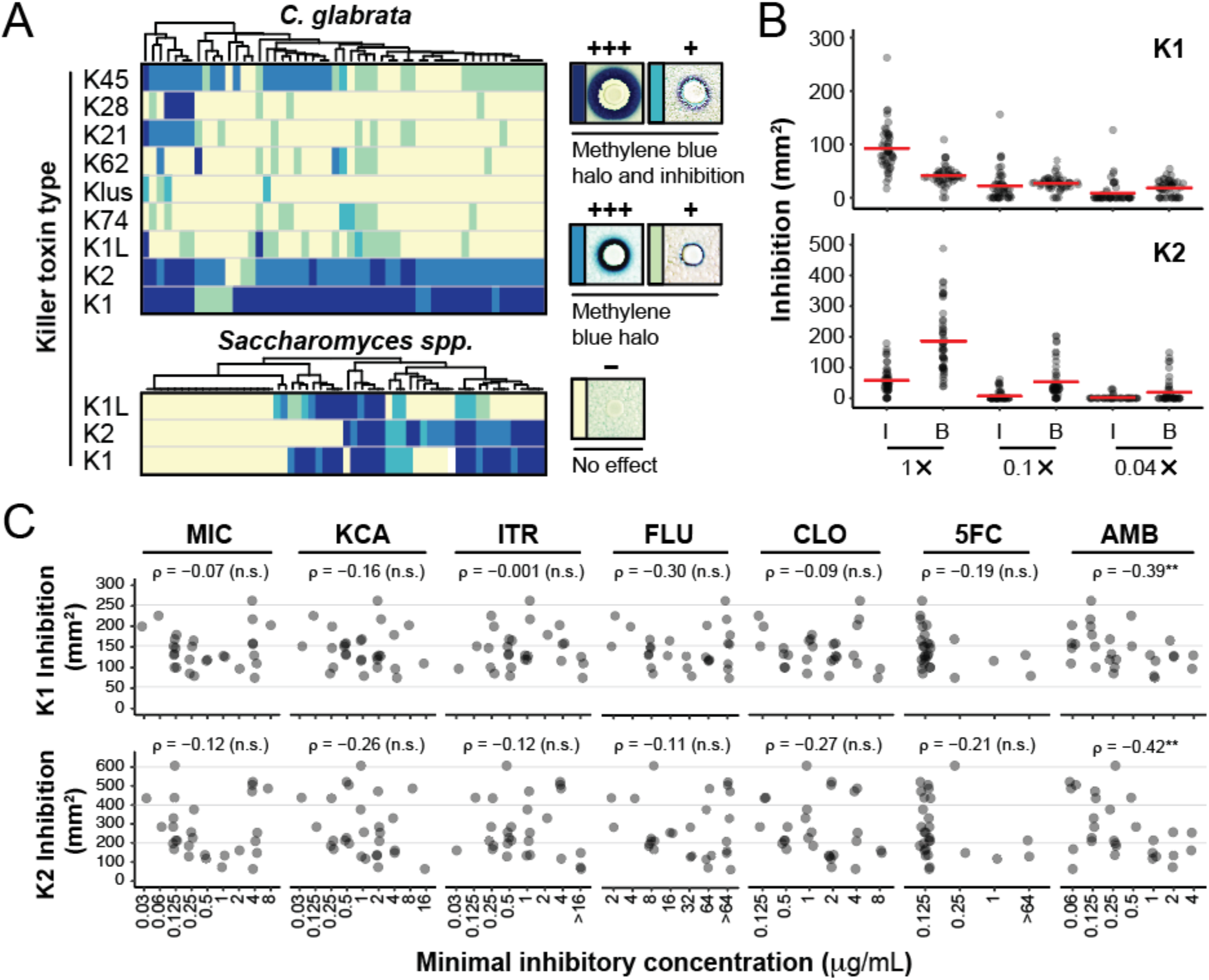
Drug resistant clinical isolates of *C. glabrata* from the vagina are most susceptible to the ionophoric killer toxins K1 and K2. (A) Cluster analysis of the susceptibility of 53 isolates of *C. glabrata* (top) and 53 strains of *Saccharomyces* species (bottom) to different types of killer yeasts as assayed on agar plates. (B) The susceptibility of 50 isolates of *C. glabrata* to partially purified K1 and K2 killer toxins showing the mean killer toxin activity based on the area of complete growth inhibition (I) or methylene blue staining (B) on agar. The 1× concentration of K1 used was 13 g/mL as measured by Western blotting using a custom antibody raised to a K1 derived peptide. (C) The minimum inhibitory concentration of seven antifungal drugs against 27 Wayne State clinical isolates of *C. glabrata* compared to the total area (methylene blue staining and zone of growth inhibition) of cytotoxicity caused by K1 or K2 killer toxins. Correlation and significance values were calculated by Spearman’s rank correlation analysis (n.s. = not significant, **p<0.05). Antifungal drugs assayed: fluconazole (FLU), clotrimazole (CLO), ketoconazole (KCA), miconazole (MIC), itraconazole (ITR), amphotericin B (AMB), and flucytosine (5FC).

Clinical isolates of *C. glabrata* were found to vary in their resistance to antifungals used to treat both VVC and invasive candidemia using the NCCLS M27-A method to calculate MIC values (Fig. 2C) and the disk diffusion assay (Fig. S2). Even when *C. glabrata* isolates were highly resistant to clinical antifungals they remained susceptible to acute K1 and K2 exposure. There was no significant correlation between drug resistance and killer toxin susceptibility (Fig. 2C and S2), except a weak correlation between K1 and K2 resistance and amphotericin B resistance (p<0.05) (Fig. 2C and S2).

Killer toxins are notoriously strain- and species-specific to the point that they have been used to identify different strains of pathogenic yeasts (26). However, the data presented in this study show that most types of known *Saccharomyces* killer yeasts can inhibit the growth of *C*. *glabrata.* Specifically, the qualitative screening of killer yeasts identified that the ionophoric toxins K1 and K2 were broadly inhibitory to vaginal isolates of *C*. *glabrata* and that purified toxins inhibited growth in a concentration-dependent manner. The *C. glabrata* cell wall is structurally similar to *S. cerevisiae* and this would suggest that K1 and K2 bind the *C. glabrata* cell wall β-1,6 glucan as the primary receptor (27). *C. glabrata* also expresses a homolog of *S. cerevisiae* Kre1p, which is the GPI-anchored secondary membrane receptor used by both K1 and K2 that enables membrane attack (28). The mechanism of *C. glabrata* intoxication is likely to involve the disruption of ion homeostasis by pore formation in the plasma membrane, as has been shown for *S. cerevisiae* (29). However, cell wall binding and the presence of Kre1p are not sufficient for intoxication, as K1 is able to bind the cell wall and utilize Kre1p of *C. albicans*, which is intrinsically K1 resistant (30, 31). Furthermore, it is unclear why other species of the *Nakaseomyces* genus (*C. nivariensis* and *C. bracarensis*) that are closely related to *C. glabrata* are more resistant to inhibition by the same killer yeasts.

The alteration of ergosterol biosynthesis can cause resistance to azoles and amphotericin B in *Candida* yeasts (32–35), and we find that the latter significantly correlates with increased K1 and K2 resistance (Fig. 2C). As amphotericin resistance can be caused by a reduction in the concentration of membrane ergosterol, increased K1 and K2 resistance in *C. glabrata* could be due to alterations in the composition, fluidity, and permeability of the yeast plasma membrane. Similar protection from K1 intoxication is observed in strains of *S. cerevisiae* with defects in ergosterol biosynthesis (36). Natural K1 resistant strains of *S. cerevisiae* also have a lower expression of ergosterol biosynthesis genes and a reduced concentration of ergosterol esters (37). The depletion of membrane sterols can result in resistance to many cytotoxic proteins (38–42). Moreover, sterol-rich membrane microdomains (lipid rafts) provide a nucleation point for protein toxins that bind raft-localized receptors (43–45). The depletion or redistribution of cholesterol can disrupt raft integrity and inhibit the binding of toxins to their cognate receptor (43, 46). Thus, susceptibility of *C. glabrata* to K1 and K2 could be influenced by membrane ergosterol and the localization and function of the GPI-anchored secondary membrane receptor, Kre1p.

The screening of killer yeasts has served to identify the unique activity of K1 and K2 against *C. glabrata* suggesting that they could be useful as novel antifungal agents. Compared to azoles, these killer toxins are fungicidal, optimally active at low pH, and are non-toxic to human cells at physiological pH(28, 47–49). Therefore, we speculate that with further mechanistic studies, formulation and stabilization, K1 and K2 could be developed for topical application to combat *C. glabrata* associated with VVC.

## Acknowledgments

The authors would like to thank Dr. Gianni Liti (University of Nice), Dr. Manfred Schmitt (Saarland University, Saarbrücken, Germany), and Dr. Reed Wickner (The National Institute of Diabetes and Digestive and Kidney Diseases) for providing strains of killer yeasts. Dr. Craig Miller (University of Idaho) is acknowledged for advice on statistical analysis. The CDC and FDA antibiotic resistance isolate bank and NRRL (ARS) culture collection provided the different species of *Candida* yeasts. The research was supported by funds provided to PAR by the Institute for Modeling Collaboration and Innovation at the University of Idaho by the National Institute of General Medical Sciences of the National Institutes of Health under award number P20GM104420, the Institutional Development Award (IDeA) from the National Institute of General Medical Sciences of the National Institutes of Health under award number P20GM103408, the National Science Foundation Cooperative Agreement DBI-0939454, and EPSCoR Track-II award number OIA1736253. Funding was also provided by the Office of Undergraduate Research at the University of Idaho (LRF, CRR, MAS, HRE, EAK). The funders had no role in study design, data collection, and analysis, decision to publish, or preparation of the manuscript and any opinions, findings, and conclusions or recommendations expressed in this material are those of the author(s) and do not necessarily reflect the views of the funders.

## Notes

### Competing Interest Statement

The authors have declared no competing interest.

### Summary of Updates

Reanalysis of the data presented in Figure 2 and the addition of a new supplementary Figure S3 showing cytotoxicity of K1 and K2 killer toxins. The discussion has been extended to relate the finding more broadly.

