## Supplementary material for "Vaginal Isolates of *Candida glabrata* are Uniquely Susceptible to Ionophoric Killer Toxins Produced by *Saccharomyces cerevisiae*": Sup. Figures

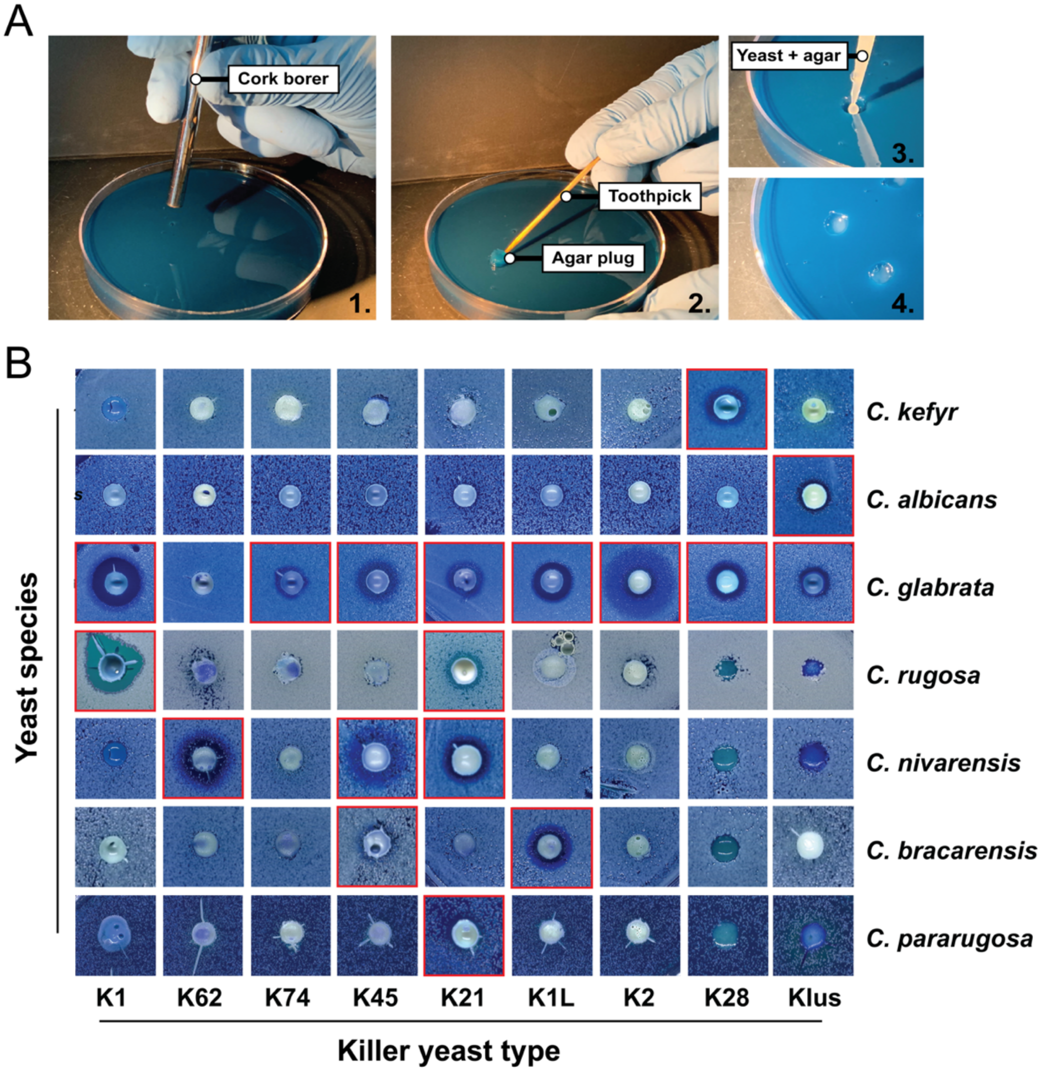


**Fig. S1. Seven species of *Candida* yeasts are inhibited by killer yeasts using a well plate assay.** (A) The killer yeast well assay method overview 1) Using a sterilized cork borer (6 mm diameter), wells were cut in an agar plate (YPD pH 4.6 with 0.003% methylene blue) that was seeded with ~1 × $10^5$ *Candida* yeast cells. 2) Wells were excavated using a sterile toothpick to remove the agar plug. 3) Approximately 5 mg of yeast from an overnight culture were mixed with cooled molten agar to a final volume of 200 $\mu$L and pipetted into each well. 4) Agar was left to set, and the plate was incubated for three days at room temperature. (B) Representative images of the inhibition of *Candida* yeasts by *Saccharomyces* killer yeasts using a well plate assay. Red highlights indicate the killer yeasts that inhibited the growth of a *Candida* yeast.

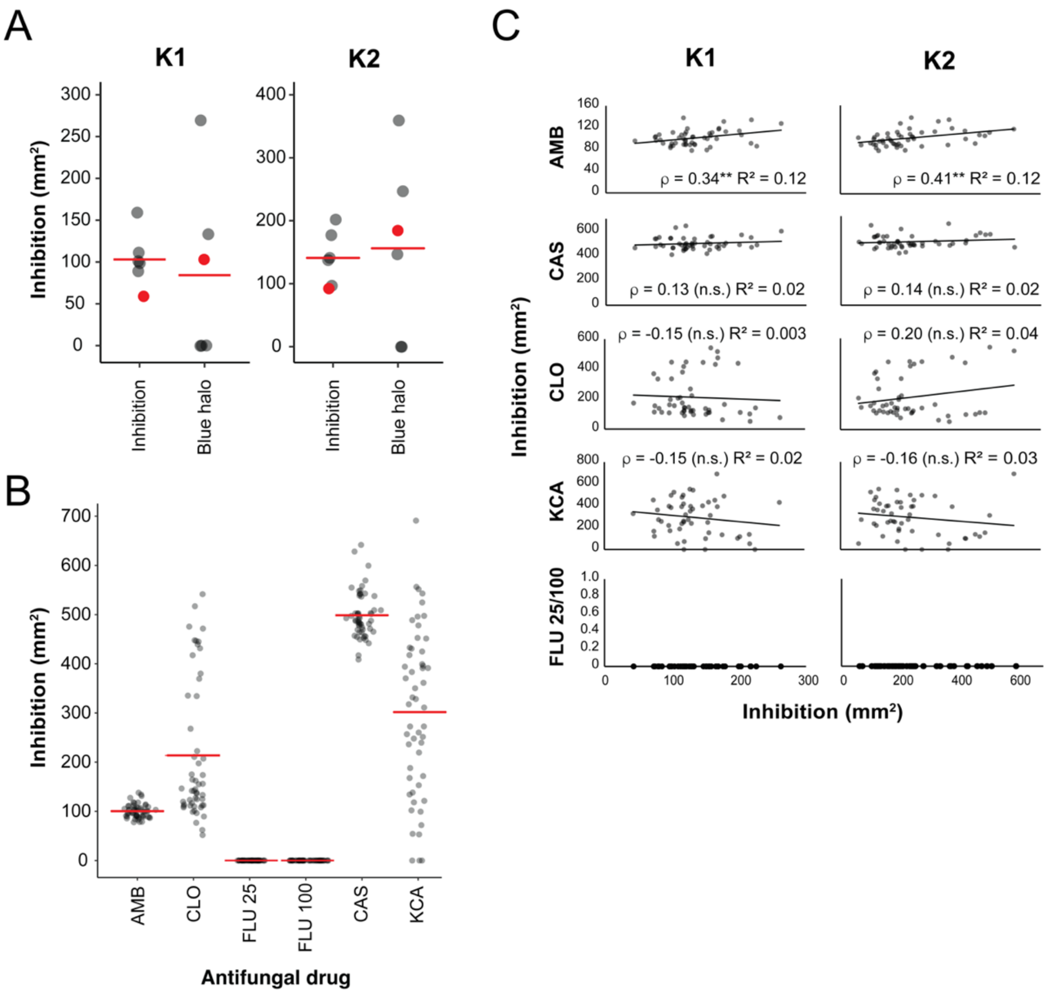


**Fig. S2. The susceptibility of *S. cerevisiae* and *C. glabrata* to killer toxins and antifungal drugs.** (A) The susceptibility of *S. cerevisiae* (grey circles) and *C. glabrata* (red circle) to partially purified K1 (13 μg/mL) and K2 killer toxins showing the mean killer toxin activity based on the area of complete growth inhibition or methylene blue staining (blue halo) on agar. (B) Areas of growth inhibition for 53 clinical isolates of *C. glabrata* by the disk diffusion assay. Horizontal red bars represent the average area of inhibition of all 53 isolates tested. (C) Spearman's correlation analysis of the relationship between the susceptibility of 50 *C. glabrata* isolates to the killer toxins K1 and K2 and clinical antifungal drugs (n.s. = not significant). All *C. glabrata* isolates were resistant to fluconazole at both 25 μg and 100 μg. Antifungal concentrations assayed: caspofungin (CAS, 5 μg), ketoconazole (KCA, 15 μg), clotrimazole (CLO, 50 μg), amphotericin B (AMB, 20 μg), and fluconazole (FLU 25, 25 μg; FLU 100, 100 μg).

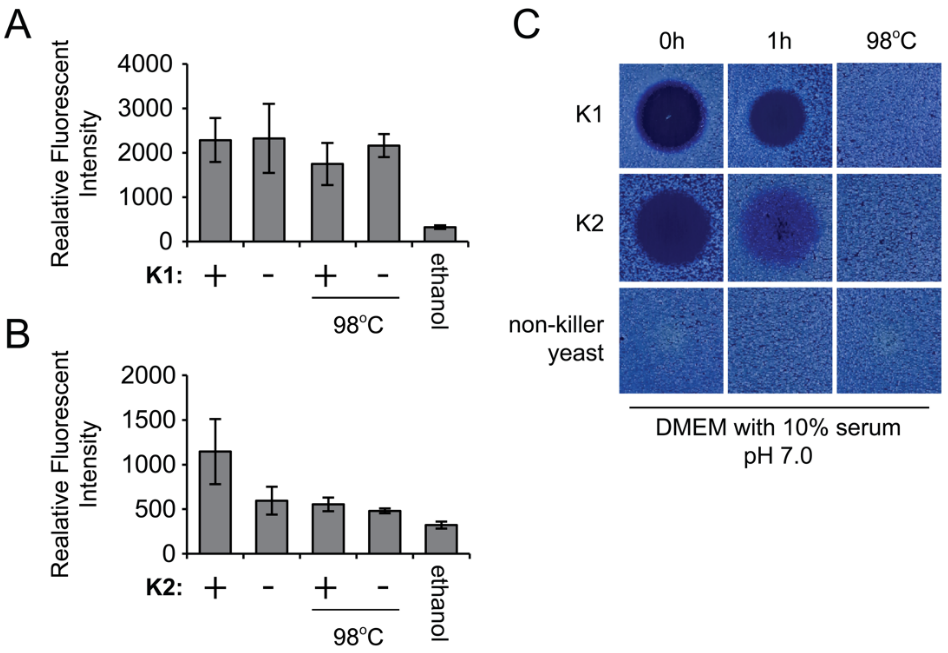


**Fig. S3. Cytotoxicity testing of K1 and K2 against cultured human epithelial cells.** Cultured HeLa cell monolayers (2 × $10^3$ cells) were exposed for 1 hour at room temperature to (A) 2.5× (0.09 μg) of ethanol precipitated K1 and (B) 2.5× of ammonium sulphate precipitated K2 suspended in DMEM with 10% serum (pH 7). Toxin preparations were aspirated from the cells and Alamar blue reagent was used to measure viability after 16 h by measuring fluorescent signal. 50% ethanol treatment was used as a control for cell death. Killer toxins heated to 98°C and precipitates from the non-killer yeast *S. cerevisiae* BY4741 were used as no toxin control samples (-). All data points represent an average of three independent repeats with standard deviation. (C) Killer toxin preparations suspended in DMEM with 10% serum (pH 7) were assayed for activity against *C. glabrata* before (0h) and after (1h) exposure to human cells, and after heating to 98°C.

| Species | Strain | *S. cerevisiae* | | | | *S. paradoxus* | | | | |
|---|---|---|---|---|---|---|---|---|---|---|
| | | CYC1058 | BJH001 | MS300c | DSM70459 | OS169 | Y8.5 | OS78 | OS40 | Y-63717 |
| *C. tropicalis* | AR 0345 | N | N | N | N | N | N | N | N | N |
| *C. parapsilosis* | AR 0342 | N | N | N | N | N | N | N | N | N |
| *C. auris* | AR 0388 | N | N | N | N | N | N | N | N | N |
| *C. duabushaemulionii* | AR 0392 | N | N | N | N | N | N | N | N | N |
| *C. krusei* | AR 0393 | N | N | N | N | N | N | N | N | N |
| *C. pelliculosa* | AR 0586 | N | N | N | N | N | N | N | N | N |
| *C. pararugosa* | AR 0587 | N | N | N | N | N | N | N | Y | N |
| *C. kefyr* | AR 0588 | N | N | Y | N | N | N | N | N | N |
| *C. guilliermondii* | AR 0590 | N | N | N | N | N | N | N | N | N |
| *C. glabrata* | ATCC2001 | Y | Y | Y | Y | N | Y | Y | Y | Y |
| *C. railenensis* | RCaA001 | N | N | N | N | N | N | N | N | N |
| *C. rugosa* | NCYC2726 | N | Y | N | N | N | N | N | Y | N |
| *C. tropicalis* | NCYC2423 | N | N | N | N | N | N | N | N | N |
| *C. nivarensis* | Y-48269 | N | N | N | N | Y | N | Y | Y | N |
| *C. bracarensis* | Y-48270 | N | N | N | N | N | N | Y | N | Y |
| *C. albicans* | Y-1289C | N | N | N | Y | N | N | N | N | N |
| **Killer toxin type** | | **K2** | **K1** | **K28** | **Klus** | **K62** | **K74** | **K45** | **K21** | **K1L** |

**Table S1. A summary of the killer toxin susceptibility of 16 species of *Candida* yeasts using a well plate assay.** Growth inhibition was characterized as an absence of yeast growth or zones of methylene blue staining. Evidence of any growth inhibition is indicated by 'Y' whereas no visible growth inhibition is represented by 'N'. Species of yeasts that were inhibited by killer toxins were retested at least once to confirm susceptibility to a killer toxin (n = 2).
